# Isolation of the murine Glut1 deficient thalamocortical circuit: wavelet characterization and reverse glucose dependence of low and gamma frequency oscillations

**DOI:** 10.1101/2023.06.05.543611

**Authors:** Elysandra M. Solis, Levi B. Good, Rafael Granja Vázquez, Sourav Patnaik, Ana G. Hernandez-Reynoso, Qian Ma, Gustavo Angulo, Aksharkumar Dobariya, Stuart F. Cogan, Joseph J. Pancrazio, Juan M. Pascual, Vikram Jakkamsetti

**Affiliations:** Department of Bioengineering; The University of Texas at Dallas, Richardson, Texas, USA; Rare Brain Disorders Program; The University of Texas Southwestern Medical Center, Dallas, Texas, USA; Department of Neurology; The University of Texas Southwestern Medical Center, Dallas, Texas, USA; Department of Physiology; The University of Texas Southwestern Medical Center, Dallas, Texas, USA; Department of Pediatrics; The University of Texas Southwestern Medical Center, Dallas, Texas, USA; Eugene McDermott Center for Human Growth & Development / Center for Human Genetics; The University of Texas Southwestern Medical Center, Dallas, Texas, USA

**Keywords:** Glucose transporter, Glut1, metabolism, thalamus, multielectrode array, oscillations

## Abstract

Glucose represents the principal brain energy source. Thus, not unexpectedly, genetic glucose transporter 1 (Glut1) deficiency (G1D) manifests with encephalopathy. G1D seizures, which constitute a prominent disease manifestation, often prove refractory to medications but may respond to therapeutic diets. These seizures are associated with aberrant thalamocortical oscillations as inferred from human electroencephalography and functional imaging. Mouse electrophysiological recordings indicate that inhibitory neuron failure in thalamus and cortex underlies these abnormalities. This provides the motivation to develop a neural circuit testbed to characterize the mechanisms of thalamocortical synchronization and the effects of known or novel interventions. To this end, we used mouse thalamocortical slices on multielectrode arrays and characterized spontaneous low frequency oscillations and less frequent 30-50 Hz or gamma oscillations under near-physiological bath glucose concentration. Using the cortical recordings from layer IV, we quantified oscillation epochs via an automated wavelet-based algorithm. This method proved analytically superior to power spectral density, short-time Fourier transform or amplitude-threshold detection. As expected from human observations, increased bath glucose reduced the lower frequency oscillations while augmenting the gamma oscillations, likely reflecting strengthened inhibitory neuron activity. This approach provides an *ex vivo* method for the evaluation of mechanisms, fuels, and pharmacological agents in a crucial G1D epileptogenic circuit.

## 1. Introduction

The most common form of glucose transporter type I deficiency (Glut1 deficiency, G1D), a metabolic encephalopathy associated with mutations of the *SLC2A1* gene, constitutes a form of medication-refractory epilepsy (Pascual and Ronen, 2015). Glut1 transporters reside in red blood cells, cells of the blood brain barrier and astrocyte membranes, all of which participate in the influx of glucose into and through the brain (Wang et al., 2023). However, elimination of the blood brain barrier in mouse brain slices, with the consequent abolition of red blood cell and capillary barrier influences, does not abolish hyperexcitability (Marin-Valencia et al., 2012). This suggests that important determinants of abnormal excitability in G1D reside within the rest of the cellular matrix of neural tissue.

Several special aspects of G1D impact the mechanistic study of such determinants and their potential therapeutic modulation. First, only astrocytes are deficient in Glut1, since neurons primarily express Glut3, but it is neurons that ultimately give rise to cerebral hyperexcitability or seizures. The simplest interpretation of this phenomenon is that the deficient transfer of astrocyte metabolites or signaling is central to the disorder. This hyperexcitability is unexpected considering the reduction in neural fuel state that characterizes G1D, which may rather be expected to lead to hypoexcitability. This apparent paradox is also noted in other energy metabolism defects (Jakkamsetti et al., 2019). Thus, studies in neural tissue rather than isolated cells are essential to preserve astrocyte-neuron relations. Second, observational behavioral studies in rodents are mechanistically limited by virtue of the predominance of unnoticeable, absence-like seizures in the disorder (Marin-Valencia et al., 2012). Many of these seizures are similarly visibly undetectable in G1D persons (Rajasekaran et al., 2022b). Third, seizures are brain fuel responsive, including dietary therapeutic sources of acetyl coenzyme A or succinyl coenzyme A (Marin-Valencia et al., 2013;Pascual et al., 2014;Rajasekaran et al., 2022b).

Fourth, human G1D seizure genesis is centered on the thalamocortical circuit but rapidly propagates to the rest of the brain (Rajasekaran et al., 2022b), such that electroencephalographic recordings (EEG) reveal apparently generalized onset epilepsy. This limits the value of EEG to elucidate mechanisms without resorting to source localization estimates (Rajasekaran et al., 2022b). Fifth, despite a traditional paucity of glucose transporter activators, an array of commercialized drugs and other poorly biologically characterized compounds robustly enhance glucose transport into cancer cells rich in Glut1 (Kathote et al., 2023), enabling the investigation of glucose influx-enhancing therapies. These considerations provide the motivation to develop an *ex vivo* circuit preparation that faithfully replicates a known neurophysiological G1D phenotype as a testbed for investigating treatments.

The thalamocortical circuit constitutes such an epileptogenesis substrate in G1D. Work by our group and others has explored the thalamocortical circuit in humans in relation to G1D seizures (Rajasekaran et al., 2022b). Notably, simultaneous functional magnetic resonance imaging and electroencephalography (EEG) revealed that human ictal activation occurs in several brain regions including the thalamus and somatosensory cortex. The electrographic correlate of this activation is a generalized, variable-duration 2-4 Hz oscillatory discharge which can be precipitated by fasting. Nonlinear source localization analysis of the discharge indicated an origin in the thalamocortical area, which also proved abnormal by fluorodeoxyglucose positron emission tomography. Synaptic function in mouse coronal brain slices revealed disinhibition of cortical pyramidal neurons and decreased synaptic inhibition of thalamic relay cells. This disinhibition was reversed by neural carbon sources such as glucose or acetate (Rajasekaran et al., 2022b). Together, these findings indicated that depletion of cerebral glucose flux impairs synaptic inhibition in the thalamocortical circuit and gives rise to aberrant thalamocortical oscillations. This explains the apparent hyperexcitability paradox cited above. Consistent with this interpretation, thalamocortical slices obtained from G1D mice displayed spontaneous electrical oscillations absent in control mouse slices (Marin-Valencia et al., 2012).

These considerations led us to develop a thalamocortical slice recording and analysis method capable of reflecting its modulation by energy metabolism after modification of bath glucose concentration. We reasoned that the characterization of thalamocortical oscillations, including those infrequent or brief, and their modulation by glucose should facilitate the investigation of epileptogenesis mechanisms and may even enable the high-throughput interrogation of metabolic fuels and drugs in a key G1D circuit. To enable quantification of low amplitude electrophysiologic oscillatory activity, we demonstrate a wavelet analysis approach which, based on simulated data, offers accurate detection from low signal-to-noise recordings. We also demonstrate, for the first time, that gamma activity can be enhanced by brain fuel in G1D. Overall, our work describes a methodological approach for investigation of epileptogenic mechanisms or treatment interventions.

## 2. Materials and methods

This study was approved by the Institutional Animal Care and Use Committee of The University of Texas Southwestern Medical Center and of The University of Texas at Dallas. All other relevant institutional regulations were followed, in addition to ARRIVE (Animal research: reporting of *in vivo* experiments) guidelines.

### 2.1. Mouse model

We used approximately equal male and female numbers of stable antisense G1D mice (Marin-Valencia et al., 2012;Rajasekaran et al., 2022b). This model was chosen because it displays a reduction in cerebral Glut1 comparable to the effect of some heterologously expressed human mutations and to human G1D red blood cells (Pascual et al., 2008). This is associated with reduced cerebral acetyl coenzyme A content, frequent electrical oscillations in thalamocortical brain slices under reduced glucose, electrographic seizures and profound ataxia (Marin-Valencia et al., 2012;Marin-Valencia et al., 2013).

### 2.2. Thalamocortical slice preparation

G1D mice aged 28 to 35 days were anesthetized with isoflurane prior to brief systemic perfusion with artificial cerebrospinal fluid (ACSF) containing (in mM) 93 N-methyl-d-glucamine, 2.5 KCl, 1.2 NaH_2_PO_4_, 30 NaHCO_3_, 20 HEPES, 10 glucose, 5 MgSO_4_, 0.5 CaCl_2_ and saturated with 95% O_2_-5%CO_2_. The mice were decapitated and the brains extracted and immediately immersed in ice cold ACSF. Sagittal-cut brains were mounted on a vibratome stage (Leica VT1000S) and 350 μm thick thalamocortical slices sectioned at a 55° angle (**Figure 1A** and **B**) generated slices as described (Agmon and Connors, 1991) with some modifications to allow contribution of slices from both hemispheres. In essence, the slicing was done at a 55° angle from the horizontal plane of the brain (in other words at a 35 degree angle from the vertical axis and parallel to the barrels (Furuta et al., 2009;Liew et al., 2021)). Since a sagittal cut was made prior to slicing, vibratome mediated slicing released two potentially usable slices, one from each hemisphere.

**Figure 1.**
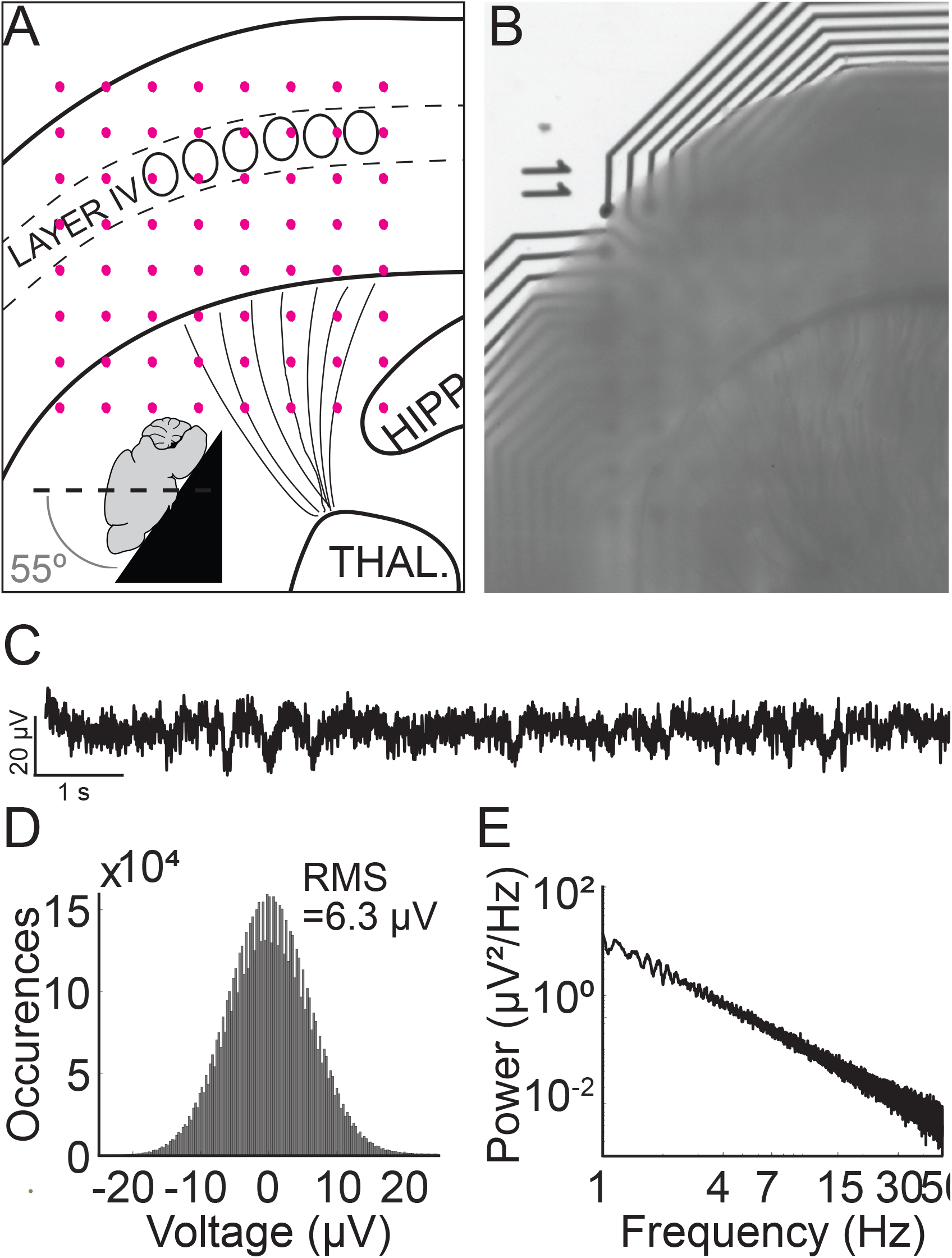
Thalamocortical slice preparation and electrophysiological properties. A) Schematic mouse thalamocortical slice illustrating cortex layer IV somatosensory barrels. Pink dots represent the underlying 8 × 8 electrode grid. The inset schema depicts the slicing angle. B) Thalamocortical slice preparation over an 8 × 8 recording electrode grid. “11” denotes the position of the first row and first column electrode. C) Recorded trace from a G1D mouse slice bathed in 2.5 mM glucose from an electrode in the 8 × 8 grid that corresponds to layer IV barrel cortex. A brief ∼ 1 second, 2 Hz oscillation at about 2 seconds from the onset of the recording trace is apparent. D) Histogram of a 10-minute trace shows the Gaussian distribution of overt background activity with a root-mean-square value of 6.3 μV. E) Power spectral density of a 10-minute recorded trace from layer IV barrel cortex. Note 1/f scaling and lack of any prominent peak corresponding to infrequent oscillation epochs.

Slices were maintained in continuously carbogenated (95% O_2_ and 5% CO_2_) ACSF containing (in mM) 118 NaCl, 2.5 KCl, 1.25 NaH_2_PO_4_, 26 NaHCO_3_, 10 glucose, 2 MgSO_4_, 2 CaCl_2_ in a 34°C water bath for 20 min and then maintained at room temperature for 40 min. Slices were then observed under the microscope (Leica M205FA) to ensure that thalamocortical connections and barrel cortex structures were preserved and aligned. The final decision on including a slice recording for analysis was made after ensuring a) the slice had clearly visible barrels in layer 4 under the microscope b) thalamocortical radiations depicting thalamocortical connections were distinctly visible c) the slice appeared healthy and viable in its electrophysiological recordings. i.e. the recording was not flat with absence of undulations and did not have an absent 1/f power spectral density of the baseline that would be indicative of non-living brain tissue d) the orientation of the recording grid adequately covered at least all cortical layers and did not visibly move over time.

Only 2 or 3 slices per brain obtained in this manner were deemed suitable for experimentation or subsequent analysis.

### 2.3. Electrophysiological recording

#### 2.3.1. Acute slice recordings

Each thalamocortical slice was positioned under optical magnification inside a well of a multi-well multielectrode array (MEA) system such that the cerebral cortex of the slice interfaced with an 8 × 8 square electrode grid (M384-tMEA-6W, Axion Biosystems) (**Figure 1A** and **B**). Electrophysiological recordings utilized an Axion Maestro Classic system (Axion Biosystems), sampled at 12.5 kHz and band pass filtered at 0.1 – 300 Hz (a frequency range used for local field potential (LFP) assay). AxIS software (Axion Biosystems) ensured the slices were maintained at 32°C while providing continuous carbogen infusion. The first recording solution consisted of low-glucose ACSF containing (in mM) 118 NaCl, 2.5 KCl, 1.25 NaH_2_PO_4_, 26 NaHCO_3_, 2.5 glucose, 2 MgSO_4_, 2 CaCl_2_. Media exchange was manually carried out between this first and the second (high glucose) ACSF solution using a pipette. To this end, the solution containing 2.5 mM glucose for baseline recordings was replaced manually with a solution containing 20 mM glucose. We did not adjust the osmolarity when switching from the ACSF containing 2.5 mM to that containing 20 mM glucose, because below a concentration of 26 mM, glucose in brain slices does not exert an osmotic effect impacting excitability (Ballyk et al., 1991). This is likely due to glucose uptake and utilization by Glut1, Glut3 and Glut6 receptors in the rodent brain as compared to impermeant compounds like mannitol which do exert an osmotic effect(Ballyk et al., 1991). A single pulse (cathodic, biphasic, 100 ms duration) was provided to the slice via a single electrode within the MEA grid utilizing the e-Stim suite (AxIS, Axion Biosystems) to interrogate slice viability. The testing sequence consisted of modulating the concentration of glucose in the ACSF (2.5 mM glucose for 30 minutes followed by 20 mM glucose concentration for 30 minutes) and recording the electrical activity over the course of 3 × 10 minute segments. This approach yielded consistent slice viability data from 6 slices Glut1D mice (2 each from 3 Glut1D mice) and 7 slices from control mice (1,3,1,2 slices from 4 wild-type mice).

#### 2.3.2. Simulated signals

A simulated time-series signal with similar sampling frequency (12.5 kHz) and duration (10 minutes) was generated to compare the ability to detect simulated oscillation epochs by different algorithms. A 1/f baseline noise signal was chosen because brain local field potentials, including our recorded signals follow a 1/f scaling at lower frequencies (Bédard and Destexhe, 2009) (**Figure 1C-E**). First, a simulated 1/f baseline noise signal was generated. This was achieved by designing a finite impulse response (FIR) filter with a ‘1/f’ passband and then filtering Gaussian random noise through it. The root-mean-square (RMS) of this signal represented baseline noise. Separately, a two second simulated epoch oscillating at 3 Hz was generated and its amplitude adjusted to twice the RMS value of baseline noise (**Figure 2A**). This two-second simulated epoch was added 20 times at random time-points to the simulated baseline noise to simulate 20 epochs of 3 Hz oscillation occurring randomly over 10 minutes of recording. This was repeated with the following amplitude adjustments for the simulated epoch: 2 ×, 1 ×, 0.5 ×. 0.25 ×, 0.125 ×, 0.0625 ×, 0.03125 × RMS noise (**Figure 2A**). Of note, “baseline noise” refers to the RMS of a 600 second recorded or simulated signal. Since this signal includes outlier deflections, any expanded example using a shorter time scale will appear to exhibit less noise than the RMS value. Each simulation of a baseline was confirmed to exhibit 1/f scaling (Figure 2B) before embedding the simulated epoch.

**Figure 2.**
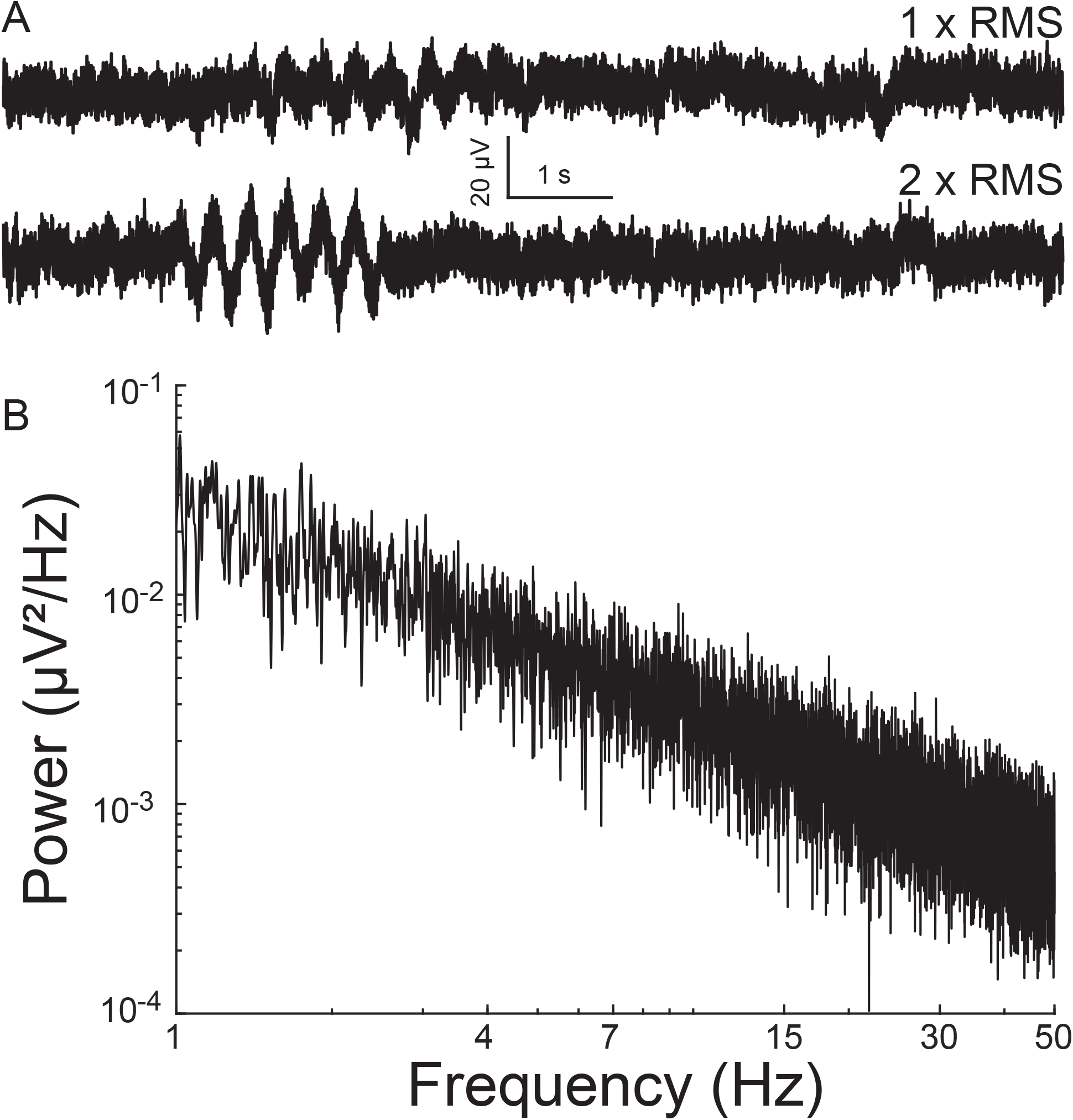
Simulated electrophysiological signal. A) Simulated 1/f baseline noise with an embedded 3 Hz oscillation epoch. The epoch amplitude is once (above) or twice (below) that of root-mean-square noise. (B) Power spectral density of baseline noise. Note 1/f scaling.

### 2.4. Data analysis

The recordings from 64 channels representing 64 electrodes were exported from the native .*raw* format to .*mat* format for analysis in MATLAB using Axion’s software tools. Each channel recording was independently analyzed.

#### Power spectrum density and short-time Fourier transform analysis

Power spectrum density curves were generated using a Fast Fourier transform for every 10 s segment of a 10 × 3 minutes (1800 seconds) recording. The average power spectra for each standard oscillatory frequency was calculated for180 segments. Short-time Fourier transform (STFT) analysis employed the “spectrogram” function offered by MATLAB. STFT analysis was conducted on each 10 s segment with a window of 0.125 seconds and a 50% overlap with consecutive windows. The window of 0.125 seconds was chosen to optimize temporal capture of oscillation epochs.

#### Wavelet-based epoch detection

A wavelet-based detection protocol for capturing oscillation. epochs was employed as previously described (Pfammatter et al., 2019). Each 30 second recorded segment (**Figure 3A** top panel, expanded for 10 seconds for ease of visualization) underwent wavelet decomposition using a *Morlet* waveform (**Figure 3A**). The analysis was sequentially conducted for each frequency bin spanning 0.3 to 64 Hz, with frequency bins set 0.25 octaves apart. The average scalogram for a frequency bin (for example from 2 to 4 Hz) was computed (**Figure 3A**). We picked 2-4 Hz as an example to describe the epoch detection algorithm since we were interested in the G1D mouse model that showed ∼ 3 Hz oscillations in a similar slice recording in our previous publication (Marin-Valencia et al., 2012). However, we did not limit our frequency range to 2-4 Hz for our simulated or acute slice recordings and provide a pan-frequency analysis for frequencies up to 64 Hz in Results. A normalized histogram of this average revealed a Gaussian distribution of points skewed to the right (**Figure 3B**), with the peak of the Gaussian distribution near zero and corresponding to the baseline, indicating that epoch activity was sparse and most points belonged to the baseline. The upper edge of the Gaussian distribution was set as the higher limit of baseline and threshold for the detection of epochs. The points above the threshold corresponded to elevations in the wavelet magnitude scalogram and this served to identify epoch events. The onset and return to baseline of the normalized wavelet magnitude marked the beginning and end of the epoch. Other than threshold, epoch events were determined to span a minimum duration of 0.5 seconds. Epoch events less than 0.5 seconds apart were considered part of a larger oscillation epoch. To clarify the presence of epoch durations shorter than the lowest frequency in our low-frequency epoch bin: We labeled low frequency epochs as those between 0.5 – 8 Hz. A complete 0.5 Hz event, for example, would be captured if it was present for > 2 second long trace, whereas a 4 Hz oscillation epoch could occur two times in a half-second long epoch. We pooled all events from 0.5 – 8 Hz epochs, thus resulting in epoch durations on average that might seem relatively too short to contain a 0.5 Hz epoch.

**Figure 3.**
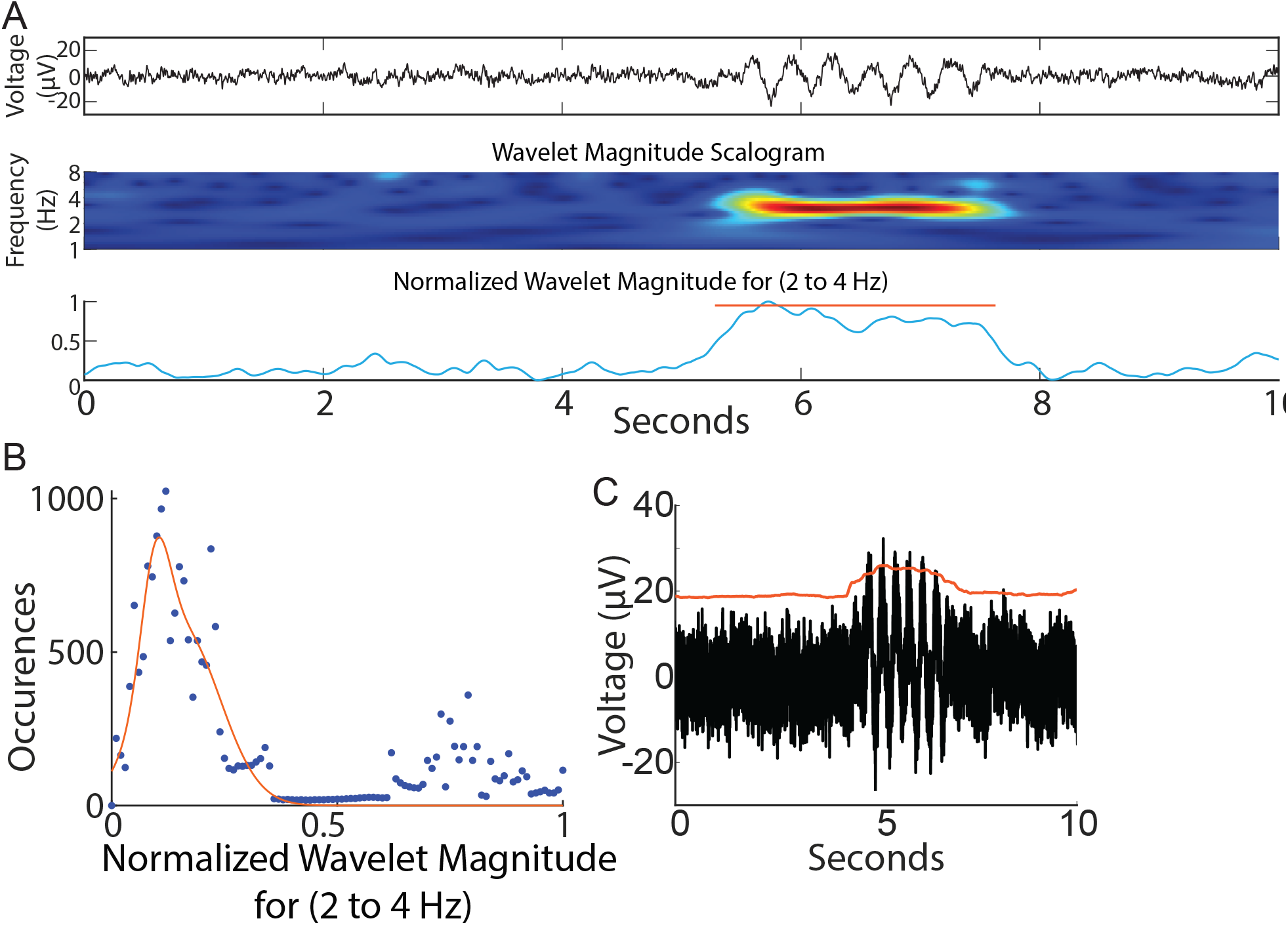
Wavelet-based and amplitude-based detection of brief oscillation epochs. A) Top panel: Simulated trace. Note brief oscillation epoch in the trace. Middle panel: Wavelet scalogram of recorded trace. Bottom panel: Average wavelet scalogram corresponding to 3 Hz (normalized) from middle panel. B) Histogram of points corresponding to normalized line in (A) bottom panel. Note majority of points measure < 0.4 units and correspond to baseline. Values greater than 0.4 units correspond to outliers above the baseline and denote the presence of an event. C) smoothed envelope (red line) captured sustained oscillations above baseline

#### Amplitude-based epoch detection

To detect oscillation epochs occurring above baseline noise, the RMS of the recording was calculated as representing baseline noise. A smoothed envelope (smoothed over 0.5 seconds, **Figure 3C**) followed the contour of baseline noise and was reliably elevated for a sustained 2-second simulated oscillation epoch (**Figure 3C**). Deflections in the envelope marked the occurrence of oscillation epochs reliably.

#### Statistical analysis

Group analysis data is depicted as mean ± standard error of mean (s.e.m.) unless otherwise noted. Statistical tests for significance are provided in the legends or the main manuscript text for the corresponding figures unless otherwise noted.. Statistical analysis was done either in MATLAB or using Graphpad Prism 9.2.

The program G*Power was employed for power analysis to determine number of samples required for statistical inference. For each analysis, Cohen’s effect size d was calculated as the absolute difference between the means divided by the pooled standard deviation. Power was set at 0.8 and α error probability was set at 0.05. Further details for each analysis is given below.

For analysis related to Figure 7D comparing WT with G1D baseline, an absolute difference of means of 6.7 (low:high frequency ratio) and pooled standard deviation of 3.69 provided an effect size of 1.82. A non-parameteric distribution of data necessitated the Wilcoxon signed rank test with a logistic distribution which (a) for a one-tailed test: provided a sample size of 5 for each group and an actual calculated power of 0.87, and (b) for a two-tailed test: provided a sample size of 6 for each group with an actual power of 0.85 as minimum required for statistical significance.

For analysis related to Figure 7D comparing G1D baseline to G1D with 20 mM glucose, an absolute difference of means of 9.52 (low:high frequency ratio) and pooled standard deviation of 4.61 provided an effect size of 2.06. With a paired-test for matched pairs with a normal distribution, (a) a one-tailed test: provided a sample size of 4 and an actual calculated power of 0.92, and (b) a two-tailed test: provided a sample size of 5 with an actual power of 0.92 as minimum required for statistical significance.

For analysis related to Figure 7E right panel comparing high frequency epochs, an absolute difference of means of 0.00125 (epochs/second) and pooled standard deviation of 0.0006 provided an effect size of 1.96. With a paired-test for matched pairs with a normal distribution, (a) a one-tailed test: provided a sample size of 4 and an actual calculated power of 0.9, and (b) a two-tailed test: provided a sample size of 5 with an actual power of 0.9 as minimum required for statistical significance.

For analysis related to Figure 7F left panel for low frequency epochs over time, an absolute difference of means of 0.008 (epochs/second) and pooled standard deviation of 0.0046 provided an effect size of 1.75. With a paired-test for matched pairs with a normal distribution, a one-tailed test: provided a sample size of 4 and an actual calculated power of 0.81, and (b) a two-tailed test: provided a sample size of 6 with an actual power of 0.9 as minimum required for statistical significance.

For analysis related to Figure 7F right panel for high frequency epochs over time (40^th^ & 60^th^ minute), an absolute difference of means of 0.0018 & 0.001 respectively (epochs/second) and pooled standard deviation of 0.001 & 0.0006 provided an effect size of -1.7 & -1.93 respectively. With a paired-test for matched pairs with a normal distribution, (a) a one-tailed test: provided a sample size of 5 & 4 and an actual calculated power of 0.9 & 0.87 respectively, and a two-tailed test: provided a sample size of 6 & 5 with an actual power of 0.88 & 0.86 as minimum required for statistical significance.

## 3. Results

### 3.1. Power spectral density analysis fails to capture infrequent events in data simulations

The power spectral density is commonly used to capture the presence of frequent oscillations in time-series signals. However, we reasoned that infrequent, low amplitude oscillations such as those typically measured in G1D slices, may not be reliably captured by this analysis. Thus, we examined the ability of a power spectrum density curve to capture infrequent oscillations using a simulated signal. For a 10-minute signal with 20 randomly inserted two-second 3 Hz oscillation epochs that were 2 times the amplitude of baseline noise, power spectral density analysis did reveal an overt or distinct peak at 3 Hz, but this peak was absent for lower amplitude oscillations (**Figure 4A**). Using recorded data, neither the baseline nor the increased-glucose condition showed any prominent frequency peak in their respective power spectral density plots (**Figure 4B**). This occurred despite the occurrence of brief oscillation epochs that could be visually confirmed in the recordings. Thus, we concluded that the infrequent and low amplitude characteristics of the oscillations contributed to their lack of detection in the power spectral density plots. This prompted exploration of alternate methods to capture individual oscillation epochs.

**Figure 4.**
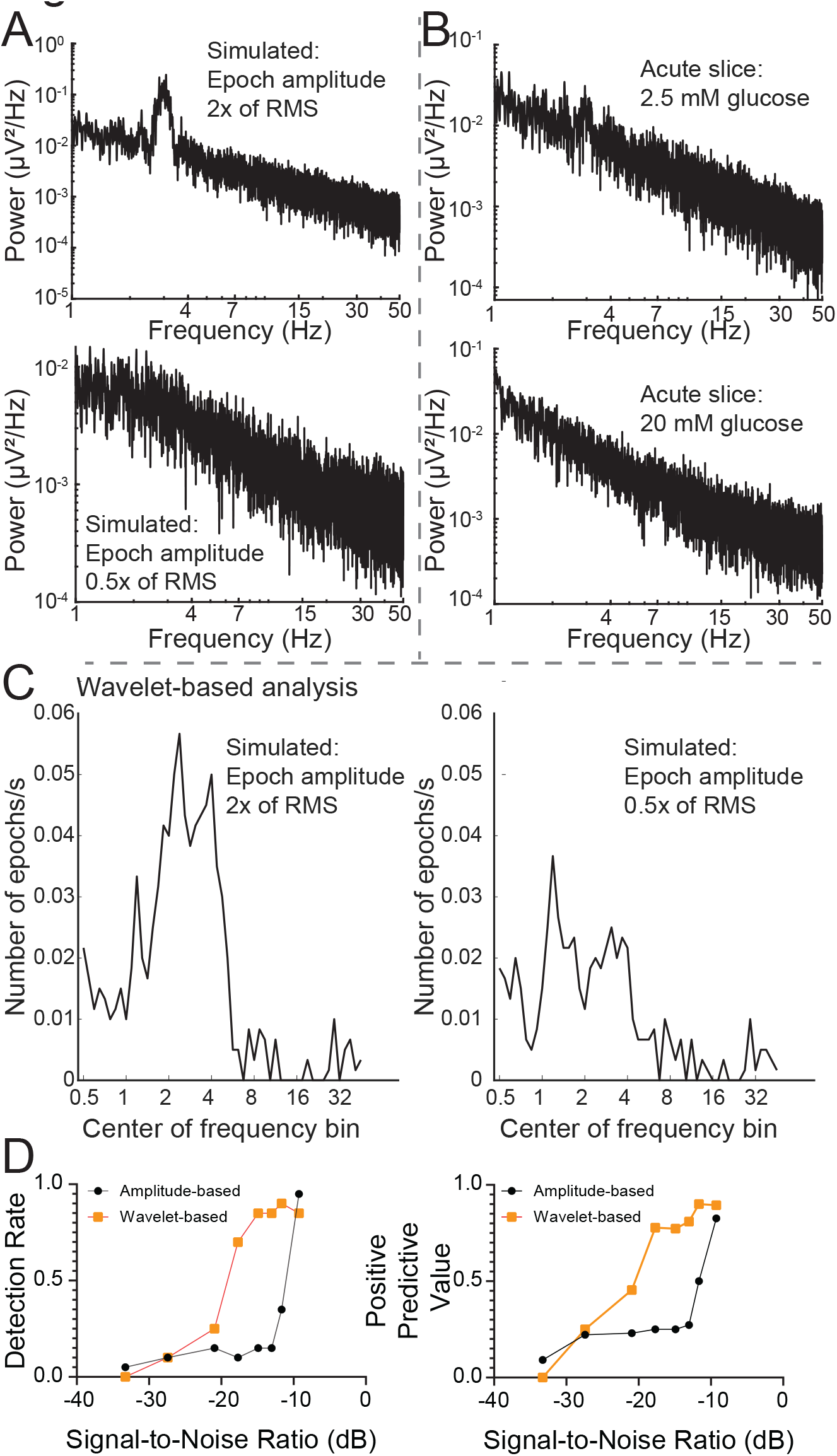
Analysis of wavelet-based algorithm performance. A) Power spectrum density plots for simulated 1/f noise with 2 × and 0.5 × RMS amplitude oscillation epochs. B) Slice data recorded at baseline and after 20 mM glucose addition. C) Wavelet-based program captured number of epochs for a 10 minute simulated recording. Note that an octave-wide frequency window was employed for the slices to allow capture of 2-4 Hz oscillations. Thus, a 3Hz oscillation epoch in simulated data will be partially captured while interrogating frequency bins an octave below (1.2 Hz) and above (5.2 Hz). D) comparison of wavelet-versus amplitude-based methods for capture of epochs

### 3.2. Wavelet based analysis reliably captures infrequent events in data simulations

We employed wavelet analysis to capture discrete epochs of oscillations in the simulated recording trace with simulated oscillation epochs. For an oscillation epoch that was twice the baseline noise (signal-to-noise ratio of -9 dB), a wavelet-based algorithm captured a 3 Hz epoch (**Figure 4C-D**). The wavelet-based approach also captured lower amplitude epochs, including some epochs with amplitudes one-half of the size of baseline noise (signal-to-noise ratio of - 20.9 dB), (**Figure 4D**). In contrast, an amplitude-based algorithm underperformed from this perspective. The algorithm detected epochs with amplitudes greater than or equal to 1.5x RMS baseline (signal-to-noise ratio of -11.7 dB), but, unlike the wavelet-based algorithm, it failed for epochs with smaller absolute amplitudes (**Figure 4D**).

### 3.3 Short-time Fourier transform based analysis has inadequate spectral resolution

Since power spectral density analysis has poor temporal resolution, we examined if short-time Fourier transform can succeed in capturing oscillation epochs. We found that STFT analysis could successfully detect simulated 3 Hz oscillation epochs in a simulated 1/f baseline noise (Figure 5A). However, temporal fidelity in capturing epochs came at the cost of frequency resolution and frequency characterization of a 3 Hz epoch was poor. For a 3 Hz epoch, our wavelet-based analysis showed a peak extending from ∼2 to ∼4 Hz, whereas the STFT based analysis showed a broader peak extending from ∼1 to ∼8 Hz (Figure 5B) indicating poor fidelity in discerning frequency of the oscillation epoch when optimized for temporal capture of these events. Hence, we utilized the wavelet-based algorithm to analyze oscillation epochs in G1D thalamocortical slices.

**Figure 5.**
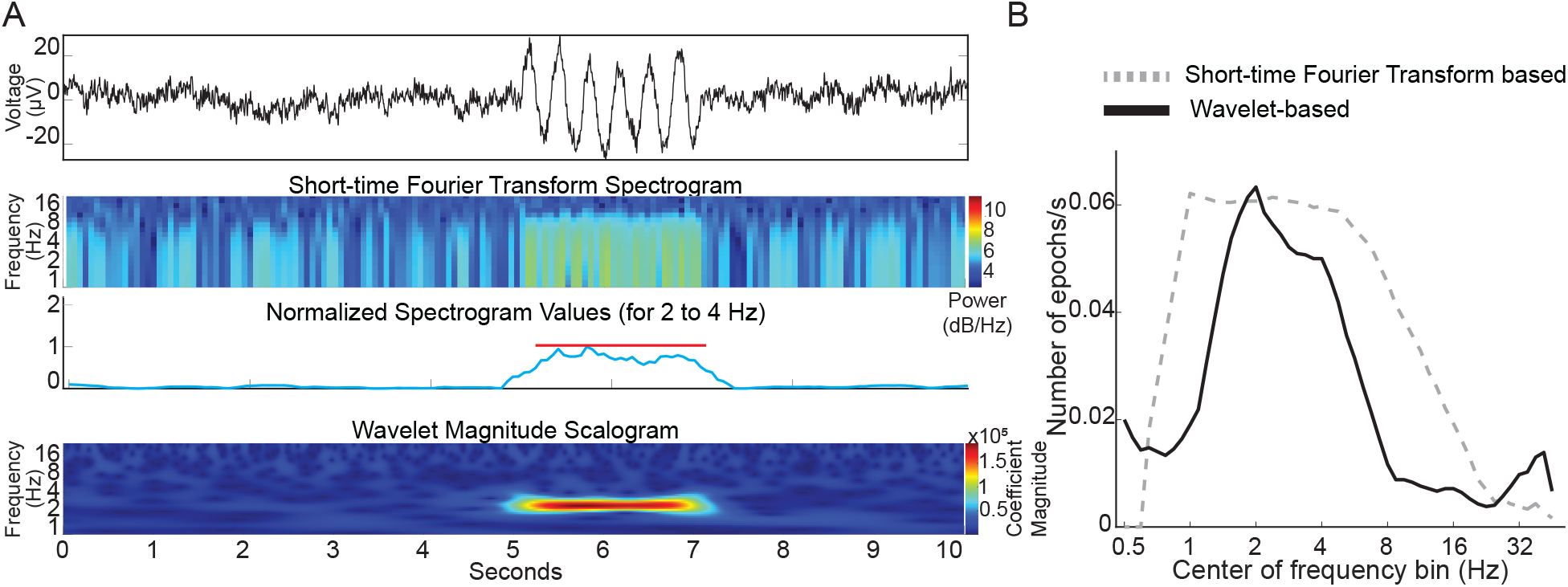
A) Top panel: Simulated trace with a brief 3 Hz oscillation epoch. Middle two panels: Short-time Fourier Transform spectrogram simulated trace with the average values corresponding to 3 Hz (normalized) below it. Lower panel: Wavelet scalogram of simulated trace. B) Comparison of STFT and wavelet-based algorithms capturing embedded 3 Hz oscillation epochs in simulated 1/f baseline noise.

### 3.4. Low and high frequency oscillation epochs

Thalamocortical slices preserved thalamocortical connections by virtue of the brain section angle. The slice was placed over an 8 × 8 electrode grid centered on the somatosensory barrel cortex and baseline recordings were acquired for 30 minutes using all 64 available channels.

We began our analysis in cortical layer IV because it is a major cortical destination of synaptic input from the thalamus. Recordings from an electrode placed against layer IV of the barrel cortex slice revealed infrequent but distinct deflections of the baseline judged as brief and distinct oscillation events (**Figure 6A**). The associated wavelet scalogram revealed the distinct presence of infrequent low frequency oscillation epochs.

**Figure 6.**
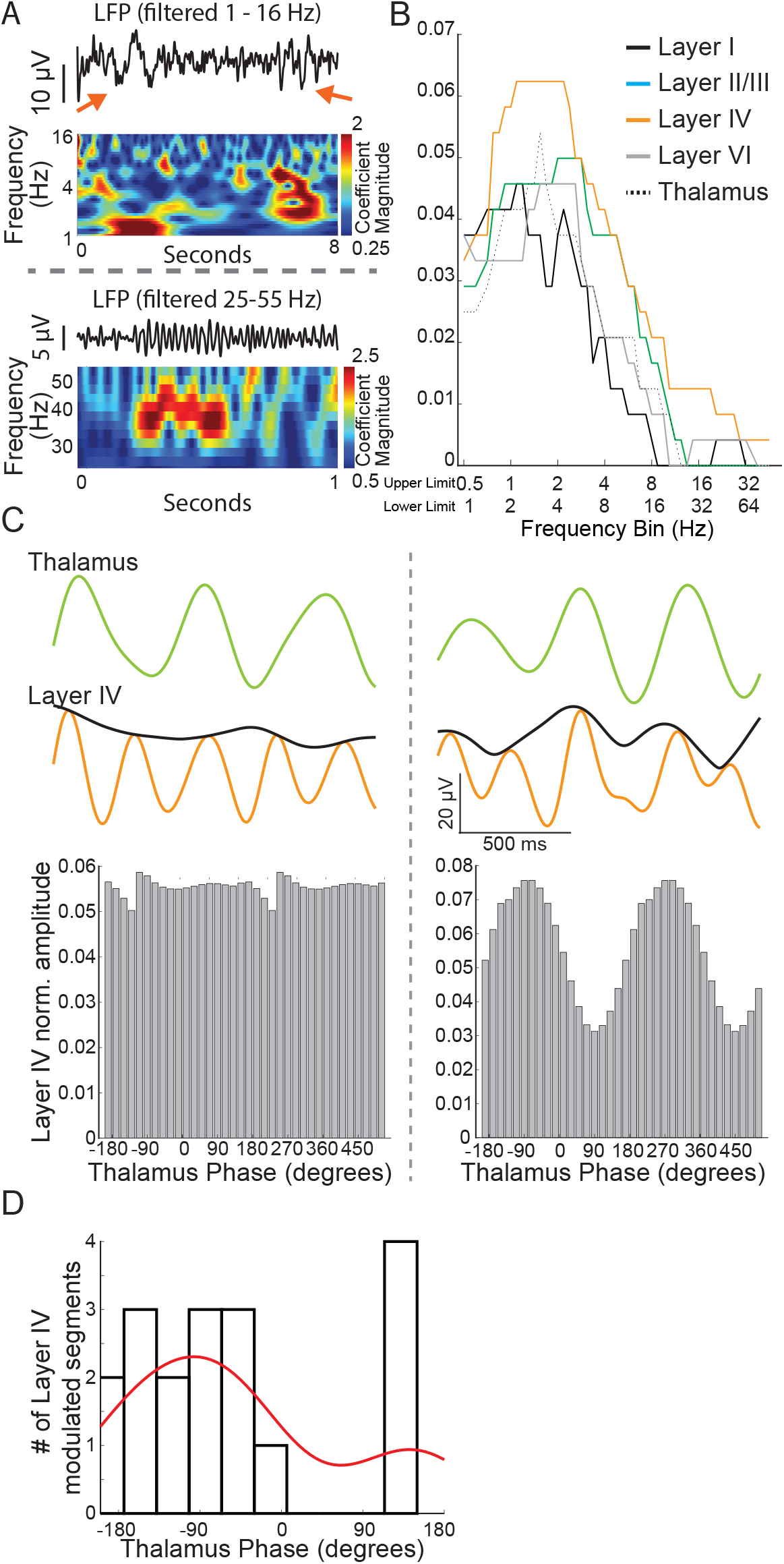
A) Recorded traces and wavelet scalogram depicting low (top panel) and high frequency (bottom panel) oscillation epochs. The red arrows in the top panel show a ∼ 1Hz (left) and ∼ 2-4 Hz oscillation epoch seen distinctly in the wavelet scalogram below. B) Number of oscillation epochs captured across different cortical regions and thalamus. Note that layer IV recordings have prominent epochs extending almost up to 4 Hz but the epochs drop off from 2 Hz for the thalamus. C) *Left panel*: a 1.5 second trace from the thalamus (filtered at 1.5 Hz) and layer IV cortex (filtered at 3 Hz) with an amplitude envelope (black color). Note lack of any impact on the cortex amplitude with thalamus activity phase. *Right panel*: Thalamus and cortex traces derived as in the left panel. Note that cortex amplitude envelope changes in tandem just before the trace derived from the thalamus. D) For most modulated segments, there was a reliable relationship (negative phase) between the phase of thalamic oscillations and the amplitude of the cortical recording.

**Figure 7.**
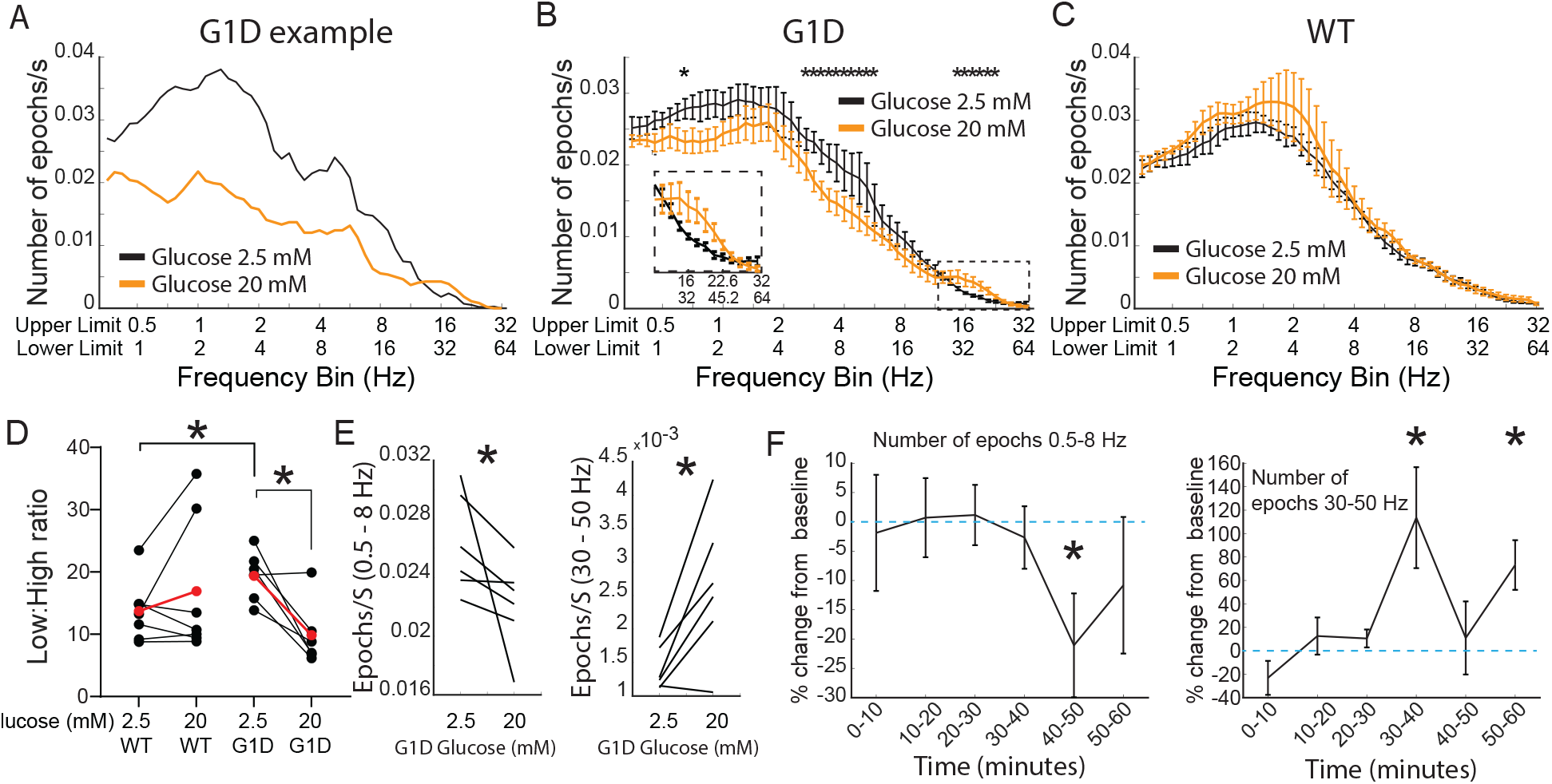
Oscillations in G1D thalamocortical slices and glucose dependence. A) Single slice modulation of oscillation epoch occurrences after addition of 20 mM glucose (orange). Black lines reflect the baseline recording (2.5 mM glucose). B-D) group data (n=6 slices, 3 G1D mice) showing modulation of oscillation epochs after 20 mM glucose (For D *=p<0.05, paired t-test for within G1D comparison, unpaired t-test with unequal variances (Welch’s test) for WT vs G1D comparison. For E *Left panel*: *=p<0.05, Wilcoxon matched-pair signed rank test; *Right panel*: *=p <0.05, paired t-test). D) Temporal profile of modulation of slices (n=6 slices, 3 G1D mice) after increased concentration of glucose. *=p<0.05,, paired t-test compared to the average epochs/sec in the 2.5 mM condition.

Of note, high-frequency local field potentials in the gamma band range require preserved energy metabolism (Kann et al., 2014) and the integrity of inhibitory neurons (Sohal et al., 2009). G1D is associated with inhibitory neuron dysfunction (Rajasekaran et al., 2022b), thus motivating us to investigate the occurrence of gamma-range oscillations and their modulation by glucose. Our wavelet-based algorithm captured these infrequent oscillations. As expected for high frequency oscillations related to elevated metabolic activity, gamma-band oscillations were brief and fewer in number than lower frequency oscillations (**Figure 6A-B**). Recordings across the cortex and thalamus confirmed that layer IV cortex had the most robust presence of ∼3 Hz oscillation epochs (Figure 6B). We saw with cross-frequency phase-amplitude coupling analysis that low frequency thalamus oscillation phases (one octave wide and centered at 1.5 Hz) have intermittent coupling with the amplitude of oscillations (one octave wide and centered at 3 Hz) in layer IV cortex, suggesting an intermittent modulatory relationship between the thalamus and pathology relevant 3 Hz oscillations in layer IV cortex (Figure 6C-D). Since layer IV recordings showed a more robust occurrence of oscillation epochs and modulatory decrease in oscillation epochs would be easier to observe in brain regions with a higher oscillation epoch count, we chose to focus our experiments on layer IV cortex.

### 3.5. Effects of glucose on low and high frequency oscillations in G1D slices

Both persons and mice with G1D exhibit neurophysiological changes upon neural fuel administration (Pascual et al., 2014;Rajasekaran et al., 2022b). Thus, we asked if our slice model captured relevant changes in low and high frequency oscillations. Specifically, human subjects with low-frequency oscillation seizure events showed a decrease in number of events with metabolic fuel and we postulated that there could be a decrease in the occurrence of low frequency oscillation epochs with addition of glucose to the slices (Rajasekaran et al., 2022b). We similarly reasoned that addition of glucose could attenuate the inhibitory neuron dysfunction characteristic of G1D and related energy metabolism defect mouse model slices (Jakkamsetti et al., 2019;Rajasekaran et al., 2022b) as reflected in a change in high frequency oscillations.

As expected, we noted that addition of elevated glucose for 30 minutes decreased low frequency oscillation occurrences and increased gamma-band frequency oscillations (**Figure 7A**). This finding was reproduced across slices, with group data indicating a significant decrease of low-frequency and increase of high-frequency oscillations (**Figure 7B**). We conducted a similar experiment in control slices (n=7 slices from 4 WT mice). When bathed in ACSF containing 2.5 mM glucose, our algorithm captured oscillation epochs in slices from WT too (**Figure 7C**). However, increasing the concentration of glucose to 20 mM did not induce any changes in the number of oscillation epochs for any frequency in slices from WT mice. Since we saw differential modulation in low compared to high frequency epochs for G1D slices, we introduced a new modulation index: a low:high frequency epoch number ratio that offers a single index to capture acute modulation of oscillation epoch activity in G1D slices when switched from 2.5 mM to 20 mM glucose (**Figure 7D**). The index showed a significant decrease with an increase in glucose concentrationin G1D slices. We found that WT slices had a lower low:high frequency ratio compared to G1D slices, suggesting an inherent oscillation frequency imbalance in G1D slices. Specifically, under low glucose, G1D slices exhibited low frequency (0.5 to 8 Hz) oscillations at a rate of 0.026 ± 0.02 per second (mean ± SEM, n=6 slices from 3 mice) with an average duration of 1.79 ± 0.03 seconds. Gamma oscillations, which were sparser, occurred at a rate of 0.0003 ± 0.0001 Hz, with an average duration of 0.96± 0.08 seconds (**Figure 7E**). Elevated glucose resulted in a significant (p <0.05, paired t-test) decrease in low frequency oscillations by 14.2± 5.9 % (or a rate of 0.0041 ± 0.0018 Hz), whereas the gamma oscillatory events rose by 86.5 ± 6 % (or a rate of 0.0012 ± 0.0005 Hz). There was no significant change in epoch duration for these low (1.75 ± 0.04 seconds) or high (1.24 ± 0.17 seconds) frequency oscillations under elevated glucose.

We also examined the temporal evolution of oscillation epoch occurrences after addition of elevated glucose (**Figure 7F**). Both low and high frequency oscillation epochs were relatively stable in number for 30 minutes for low frequency oscillations. There was no change immediately following glucose addition, indicating that solution exchange did not perturb the recording. However, 10 minutes after introduction of 20 mM glucose, there was a significant change in the occurrences of low-frequency oscillations, which decreased by 24.9 ± 7.4 % (p<0.05, paired t-test comparing average number of epochs at baseline with the relevant 10 minute time bin). Introduction of 20 mM glucose increased the frequency of high frequency oscillations for the first 10 minutes (by 115 ± 36 %, p<0.05 paired t-test compared to average epoch number at baseline) and the 20-30 the minute bin (73 ± 18 %, p<0.05, paired ttest compared to average epoch number at baseline) after increasing the concentration of glucose (**Figure 7F**).

## 4. Discussion

In this work, we have demonstrated a thalamocortical slice model that demonstrates glucose responsive electrophysiological abnormalities at low frequencies that is in congruence with glucose response ∼ 3 Hz EEG findings seen in G1D subjects. With the presence of Glut1 receptors on astrocytes, and the robust role of astrocytes in metabolism and neurotransmission relevant to glutamate and GABA (Andersen and Schousboe, 2023;Andersen et al., 2023), our study helps extend our understanding of the relationship between Glut1 deficits and downstream impact on neurotransmission and electrophysiology. We introduce an analysis approach capable of reliably quantifying associated brief extracellular potential oscillatory activity derived from MEA recordings. MEA recordings in thalamocortical slices are useful to examine the normal interplay of excitation and inhibition using electrical stimulation and pharmacological modulation (Wirth and Luscher, 2004). This approach can be expanded to study the abnormal electrical oscillations characterize several types of human epilepsy. Among these, low frequency oscillations in the 2 - 4 Hz range are typical of absence seizures (Sadleir et al., 2006). Rodent model seizures exhibit similar oscillations (Cox et al., 1997;Persad et al., 2002). In this context, several studies have employed wavelet analysis of epileptiform brain slice oscillations in various models (Jiruska et al., 2010;Gonzalez-Sulser et al., 2011).

With this precedent, we used a mouse model that recapitulates the cardinal features of human G1D including microcephaly, ataxia and epilepsy (Marin-Valencia et al., 2012). Importantly, these features are associated with reduced cortical and thalamic glucose accumulation and diminished acetyl-coenzyme A production. In contrast with other models, thalamocortical slices (which are virtually devoid of endothelial glucose flux) obtained from G1D mice display spontaneous electrical oscillations (Marin-Valencia et al., 2012). We previously characterized the cellular mechanism underlying these oscillations by assaying synaptic function in coronal brain slices. This revealed disinhibition of cortical pyramidal neurons. In contrast also with other thalamocortical epilepsies, thalamic tonic inhibition was unaltered, but synaptic inhibition of thalamic relay cells was also decreased. This disinhibition was reversed by neural carbon sources such as glucose or acetate. This allowed us to hypothesize that cerebral glucose flux regulates brain activity by modulating synaptic inhibition in the thalamocortical circuit which, in the case of G1D, may be rate-limited by glucose (Rajasekaran et al., 2022b). Thus, the resulting neurophysiological abnormalities such as disinhibition should prove amenable to increased glucose supply, as we observe.

Our findings show not only that the low frequency oscillations were attenuated with elevated glucose, but also that gamma field potential oscillations were also modulated as expected by the neurophysiological circuit correlate of depressed and then fuel-restored disinhibition. This may be due to the enhanced sensitivity of inhibitory GABAergic neurons, specifically their parvalbumin-positive subtype (or Pvalb), relative to excitatory neurons (Mann et al., 2005). We have shown that Pvalb release less neurotransmitter in mouse genetic energy metabolism failure, impacting local circuit and electrocorticographic activity (Marin-Valencia et al., 2012;Jakkamsetti et al., 2019). Pvalb activity may be more vulnerable to metabolic failure than glutamatergic neurons due to the dependence of GABA recycling on the astrocyte TCA cycle via obligatory conversion to succinate, which is not necessary for glutamate (Bak et al., 2006). We have used gamma oscillations to predict seizures in an energy metabolism human and mouse disease (Jakkamsetti et al., 2019). A consequence of these observations is that, as we have also shown in mice (Jakkamsetti et al., 2019;Jakkamsetti et al., 2022) and in persons with genetic energy metabolism disease (Pascual et al., 2014), metabolic failure in Pvalb and excitatory neurons should prove modifiable by glucose, as suggested by our results.

In contrast with our approach, notwithstanding the above references, it has been common to measure brain signal oscillations via power spectrum analysis. This involves the application of the Fast Fourier transform to the data to convert the recorded signal from the time to the frequency domain. For example, for a 10-minute recording, the power spectrum is often calculated for each 10 second recorded segment, and an average of the power spectrum across all 10 second segments in a 10-minute recording is determined. The presence of a continuous oscillation in the signal can be thus detected as a prominent increase in power for that oscillation frequency. However, this is disadvantageous for the detection of oscillations such as those common in G1D, since discontinuous, infrequent, low amplitude and brief oscillations, as previously noted in this model (Marin-Valencia et al., 2012) are likely to remain unidentified or missed. A short-time Fourier transform based analysis on the other hand, manages to temporally capture epochs, but with poor fidelity in the frequency domain. In contrast, the wavelet-based method captured brief oscillation epochs in the G1D thalamocortical slice model with adequate temporal as well as spectral fidelity. A wavelet transform highlights or facilitates the measurement of an expanded (such as a lower frequency oscillation) or compressed (such as a faster frequency oscillation) stereotypical wavelet shape at each time point. The portions of the recording that better match the expanded or compressed wavelet shape exhibit greater wavelet coefficients, thus helping identify the presence of each oscillation epoch. Thus, a wavelet transform can identify the frequency as well as the time of occurrence of infrequent oscillations. By these means, we measured low frequency oscillations in G1D slices that decreased with increased glucose. Complementarily, glucose increase was also followed by an increase in gamma oscillations.

Physiological brain fluid glucose is estimated at about 1.1 mM (McNay and Gold, 1999). Cerebrospinal (CSF) glucose concentration in our previous G1D mouse model, with the caveat that this reflects brain tissue concentration only in part (Lund-Andersen, 1979), is 1-1.5 mM (Wang et al., 2006). 2.5 mM glucose in the bath results in a concentration of ∼1.5 mM at 100 μm depth within the brain slice (Tekkok et al., 2002), which is the depth from which we estimate our recordings originate. Thus, we are recording at near physiological glucose concentration.

One of the study’s limitations is in restricting analysis to recordings from layer IV in the cortex. In this mouse model, both the cortex and thalamus exhibit inhibitory failure (Rajasekaran et al., 2022a) and both regions could contribute to the origin or sustenance of a pathological cortical recorded 3 Hz oscillation. It could be useful to know if either of the two structures was more vulnerable and initiated the 3 Hz oscillation epoch recorded in the cortex. Such an analysis would require a millisecond precision in capture of epochs since it takes ∼ 5 milliseconds for synaptic transmission from the thalamus to the cortex. Unlike capture of discrete and binary events like action potentials, detecting the exact transition of a small-amplitude natural 3 Hz oscillation to a large-amplitude pathological 3 Hz oscillation can be imprecise. More involved experiments including encouraging seizure activity with chemical agents to capture and average across multiple epochs to decrease imprecision, and separating thalamus from the cortex as has been done before in an absence seizure rodent model (D’Arcangelo et al., 2002) could help address this issue in the future. In addition, to ensure that a barrel from layer IV was recorded from, the current priority was on placement of the MEA with adequate coverage over the cortex, with occasional coverage over the thalamus. High density or custom-made MEAs with wider coverage should help simultaneous and targeted recordings from layer IV cortex and thalamus to allow a more thorough investigation of their specific involvement.

The relationship between low energy states and seizure initiation and maintenance can be complicated. In extreme low-energy states, it is not unexpected to have hyperexcitability. Low energy states will result in ATP deficits which is expected to impair NaK-ATPAse activity and glutamate handling. NaK-ATPase is one of the largest energy consumers in the brain (Engl and Attwell, 2015). Its hypofunction should impair ion balance and raise resting membrane potential, change potassium uptake by astrocytes and promote seizure initiation. Interestingly, for cells from living mice with a low-energy state mouse model (Jakkamsetti et al., 2019), there were deficits in rapid firing inhibitory neuron function and glutamate release, or deficits in inhibitory currents in a G1D mouse model (Rajasekaran et al., 2022), but resting membrane potential was normal. This suggests the possibility that NaK-ATPase fueling is prioritized at the cost of other functions during low energy states and only fails in extreme ATP deprivation. To make matters more complicated, inhibition of glycolysis by 2-deoxyglucose has been seen as both pro- and anti-convulsant in studies (Samokhina et al., 2017;Rho et al., 2019). Moreover, for seizure maintenance, it is conceivable that energy is required (Dienel et al., 2023). A lack of continued energy availability in G1D could impact continuation of seizures, thus inducing network hypoexcitability even in the presence of end-stage neuronal hyperexcitability.

## 5. Conclusions

We have developed a thalamocortical slice recording and analysis approach that can quantify extracellular local field potential oscillations related to epileptogenesis in a G1D mouse model. While the amplitude of the electrophysiological activity measured using a substrate-integrated MEA was relatively low compared to other neurophysiological activities measurable in slices, our automated wavelet-based algorithm reliably detected low SNR events as demonstrated by the data simulations. Consistent with prior work (Marin-Valencia et al., 2012), we observed well-resolved 2-4 Hz oscillations that occurred spontaneously at low bathing glucose concentrations. Increasing bath glucose concentration reduced these slow oscillation events. This contrasts with the concurrent enhancement of high frequency gamma oscillations, which are associated with local inhibition (Jakkamsetti et al., 2019;Rajasekaran et al., 2022b) (Kann et al., 2014). Our findings corroborate, at the thalamocortical circuit level, that G1D represents a metabolism-dependent disinhibited state that leads to thalamocortical oscillations.

Our slice model containing essential parts of a key neural circuit that is amenable to modulation and the method of analysis should enable the characterization of further epileptogenesis mechanisms. while enabling the screening, perhaps in high-throughput fashion after modifications (Dunlop et al., 2008), of metabolic fuels and pharmacological agents that may ultimately prove of clinical utility in the treatment of G1D. The method can be adapted to study other genetic and toxin-induced forms of abnormal thalamocortical oscillations. Specifically, the barrel somatosensory cortex, which was included in our slice preparation, and the interconnected thalamic circuitry are of interest in disorders other than G1D because they can become hypersynchronized, causing absence seizures in numerous epilepsy models (Kim et al., 1997;Huntsman et al., 1999;Macdonald et al., 2010).

## Acknowledgments

We are grateful to Gauri Kathote and Ignacio Málaga for discussions. We also thank the Glut1 Deficiency Foundation for providing a patient and scientific forum where this project was debated. All authors take responsibility for the integrity of the statements and the accuracy of the analysis. No institution participated in the design and conduct of this work; or preparation, review, or approval of the manuscript or decision to submit the manuscript for publication.

## Contribution to the field statement

Electrical oscillations, or excessive neural circuit synchronization, characterize several seizure types. Seizures in human and mouse model Glut1 deficiency are associated with oscillations between thalamus and cerebral cortex as noted by electroencephalography and functional imaging. The axons between these regions are anatomically robust, such that mouse brain sections or tissue slices can be extracted with preserved thalamocortical connections. The circuit cross-section thus obtained is viable under artificial superfusion, eliminating influences from other severed brain areas while allowing for electrophysiological recording and manipulation. Under a glucose concentration close to normal, Glut1 deficient slices display spontaneous electrical oscillations, thus allowing for the study of mechanisms and potential treatments in a manner unfeasible via other approaches. To faithfully capture the frequency and amplitude of the oscillations, we used a wavelet analysis method that enabled the identification of two types of abnormal activity: low and high, or gamma, frequency oscillations. The latter, which were less pronounced than the former, arise from inhibitory neuron activity and can suppress excess excitation. Elevated glucose, which suppresses human and mouse Glut1 deficient seizures, exerted differential effects on these oscillations as expected from a therapeutic effect, therefore highlighting the usefulness of this circuit and wavelet analysis approach.

## Conflict of interest

The authors declare that the research was conducted in the absence of any commercial or financial relationships that could be construed as a potential conflict of interest.

## Author contributions

Conception and design: JMP and JJP

Drafting of the manuscript: VJ, EMS, JMP

Experimental studies: EMS, VJ, LBG, RGV, SP, QM, GA

Data analysis: EMS, VJ, LBG, RGV, SP, AD

Critical revision of the manuscript for important intellectual content: All authors

Supervision: JMP, JJP

## Funding

This work was supported by the National Institute of Neurological Disorders and Stroke (R01-NS077015, and R01-NS078059 to JMP), by The University of Texas at Dallas Office of Research and Innovation through the Collaborative Biomedical Research Award Program (to SC and JMP) and by the Glut1 Deficiency Foundation (to JMP).

## Data availability statement

The data generated and analyzed for this study are available upon request from the corresponding author. EMS, VJ, JJP and JMP had full access to all the data in the study and take responsibility for the integrity of the data and the accuracy of the data analysis.

## Abbreviations

1/f noise: 1/frequency noise
ACSF: Artificial cerebrospinal fluid
ARRIVE: Animal research: reporting of *in vivo* experiments
EEG: electroencephalography
G1D: Glucose transporter type 1 deficiency
Glut1: Glucose transporter type 1
MEA: Multielectrode array
Pvalb: Parvalbumin-positive subtype neurons
SNR: Signal-to-noise ratio
TCA cycle: tricarboxylic acid cycle
STFT: Short-time Fourier Transform

## Notes

### Competing Interest Statement

The authors have declared no competing interest.

### Summary of Updates

This version: (1) Includes a new experiment with slices from wild-type mice (Figure 7C-D), and two new figures with analyses (2) Examines the prominence of layer IV cortex oscillations (Figure 6B) and the relationship of layer IV cortex and thalamus oscillations (Figure 6C-D) (3) Shows that Short-time Fourier Transform can detect oscillation epochs but at the cost of fidelity in the frequency domain (Figure 5A-B). (4) Includes multiple clarifications in the main text (primarily in the Methods and Discussion section) and a few in the figures/legends.

